# Mutant *C. elegans* mitofusin leads to selective removal of mtDNA heteroplasmic deletions at different rates across generations

**DOI:** 10.1101/610758

**Authors:** Lana Meshnik, Dan Bar-Yaacov, Dana Kasztan, Tali Neiger, Tal Cohen, Mor Kishner, Itay Valenci, Sara Dadon, Christopher J. Klein, Jeffery M. Vance, Yoram Nevo, Stephan Zuchner, Dan Mishmar, Anat Ben-Zvi

## Abstract

Deleterious and intact mitochondrial DNA (mtDNA) mutations frequently co-exist (heteroplasmy). Such mutations likely survive and are inherited due to complementation via the intra-cellular mitochondrial network. Hence, we hypothesized that compromised mitochondrial fusion would hamper such complementation, thereby affecting heteroplasmy inheritance. To test this hypothesis, we assessed heteroplasmy levels in three *Caenorhabditis elegans* strains carrying different heteroplasmic mtDNA deletions (ΔmtDNA) in the background of mutant mitofusin. Firstly, these animals displayed severe embryonic lethality and developmental delay. Strikingly, these phenotypes were relieved during subsequent generations in association with complete ΔmtDNA removal. Moreover, the rates of deletion loss negatively correlated with the size of mtDNA deletions, suggesting that mitochondrial fusion is essential and sensitive to the nature of the heteroplasmic mtDNA mutations. While introducing the ΔmtDNA into a *fzo-1*;*pdr-1* (PARKIN ortholog) double mutant, we observed skew in the mendelian distribution of progeny, in contrast to normal distribution in the ΔmtDNA;*fzo-1* mutant, and severely reduced brood size. Notably, the ΔmtDNA was lost across generations in association with improved phenotypes. This underlines the importance of cross-talk between mitochondrial fusion and mitophagy in modulating the inheritance of mtDNA heteroplasmy. Finally, while investigating heteroplasmy patterns in three Charcot-Marie-Tooth disease type 2A pedigrees, which carry a mutated mitofusin 2, we found a single potentially deleterious heteroplasmic mutation, whose levels were selectively reduced in the patient versus healthy maternal relatives. Taken together our findings show that when mitochondrial fusion is compromised, deleterious heteroplasmic mutations cannot evade natural selection, while inherited from generation to generation.

## Introduction

Unlike the nuclear genome, mitochondrial DNA (mtDNA) is present in multiple copies per animal cell. For instance, human somatic cells contain an average of ~1,000 mitochondria per cell, with each mitochondrion harboring 1-10 mtDNA copies [1]. Although this large intracellular mtDNA population is inherited from the maternal germline, and hence carries a single major haplotype, mtDNA molecules can differ in sequence (heteroplasmy) either due to inheritance of mutations from the ovum or due to the accumulation of changes over the lifetime of the individual [2–5]. Some of these changes may have pathological consequences [6, 7], as reflected in a variety of mitochondrial disorders, yet only upon crossing a threshold of prevalence in the cell [8]. Accordingly, the penetrance of disease-causing mutations ranges between 60-80%, depending on the symptoms and tissues that display the specific phenotype [9].

The repertoire of heteroplasmic mutations varies among cells and tissues of an individual, mainly due to replicative segregation (drift) of the mitochondria during cell division and mitochondrial bottlenecks that appear during embryo development [10]. However, it has been suggested that heteroplasmy can be modulated by non-random factors, including selection [2, 4, 5, 11]. Indeed, it has been shown that mitophagy, a mechanism of mitochondrial quality control, partially provides selection against defective mitochondria and maintains disease-causing mtDNAs below the threshold levels both in human cells [12] and in a *Caenorhabditis elegans* model [13–15]. Mitophagy requires proper fission-fusion cycles of the mitochondrial network so as to allow removal of dysfunctional mitochondria [16, 17]. In agreement with this notion, elevated heteroplasmy levels of pathological mtDNA were observed when the fission machinery was disrupted in cell culture [18]. Furthermore, reduction in heteroplasmy levels of potentially deleterious mtDNA mutations was observed when components of the fusion machinery were compromised in *Drosophila* models [19], and especially in germ cells [20]. In consistence with this notion, cell culture experiments revealed that a mixture of mtDNA molecules differing in sequence in the same cell can complement each other by the diffusion of products via the mitochondrial network, which in turn leads to a restoration of mitochondrial function [1, 14, 21]. Hence mitochondrial fusion likely allows the survival of mtDNA disease-causing mutations in cells and in turn, their transmission to the next generation [1, 8, 22]. Although these experiments suggest a molecular mechanism for the control of heteroplasmy, it remains unclear whether such mechanism also affects the transmission of heteroplasmy from generation to generation, thus potentially explaining the relatively high abundance of low-level disease-causing heteroplasmic mutations in the general population [23, 24]. We, therefore, hypothesized that interfering with the intracellular mitochondrial network by compromising the fusion machinery would hamper mitochondrial functional complementation and thus inheritance of heteroplasmic mutants.

Here, we took the first steps towards testing this hypothesis by crossing *C. elegans* harboring mitofusin (*fzo-1*) mutant to animals carrying either of three heteroplasmic mtDNA deletions, which differed in size and mtDNA positions. These experiments resulted in an embryonic lethality and developmental delay, which was alleviated in subsequent generations concomitant with a complete loss of the truncated mtDNA molecules. Since the rate of truncated mtDNA loss diverged between the heteroplasmic strains, the sensitivity of the fusion machinery to different mtDNA mutations, in addition to its interaction with mitophagy and relevance to human diseases are discussed.

## Results

### A heteroplasmic deletion reduces the fitness of *C. elegans* mitofusin (*fzo-1*) mutant

The stable heteroplasmic *C. elegans* strain *uaDf5/+* harbors a mixture of intact (+mtDNA) and ~60% of a 3.1 kb mtDNA deletion (ΔmtDNA) [25]. Although lacking four essential genes (i.e., mt-ND1, mt-ATP6, mt-ND2 and mt-Cytb), this strain displays only mild mitochondrial dysfunction [15, 25, 26]. We showed that dysfunctional PDR-1, the worm orthologue of the key mitophagy factor Parkin (PARK2), led to elevated levels of the truncated mtDNA, suggesting that mitochondrial quality control senses the presence of dysfunctional mitochondria [13]. In conjunction with this finding, RNAi knockdown of *fzo-1*, the *C. elegans* orthologue of MFN1/2, led to a slight reduction in the levels of the heteroplasmic ΔmtDNA, although without any phenotypic consequences [14]. We, therefore, asked what would be the impact of the *fzo-1(tm1133)* deletion (hereafter designated as *fzo-1(mut)*) on the inheritance of the maternal ΔmtDNA.

To this end, we crossed ΔmtDNA heteroplasmic hermaphrodites with *fzo-1(mut)* heterozygote males (Fig. 1A). After self-cross of the F1 progeny, the distribution of the genotypes in the F2 heteroplasmic progeny did not deviate from the expected Mendelian ratios, namely 26% homozygous *fzo-1(mut)*, 48.7% *fzo-1* heterozygotes (*ht*) and 25.3% *fzo-1(wt)* (p=0.96, Chi square test; Table S1). No apparent phenotypic differences were observed among F2 animals (generation 1, G1) as well as in the non-heteroplasmic *fzo-1(mut)* animals. However, we noticed that only 13±5% of the second generation (G2m; i.e., the progeny of the self-crossed *fzo-1(mut);*ΔmtDNA worms) hatched, as compared to *fzo-1(mut)* animals (67±5%; Fig. 1B). Interestingly, the 13±5% G2m animals that did hatch were developmentally delayed, and none of the animals reached adulthood after six days.

**Figure 1:**
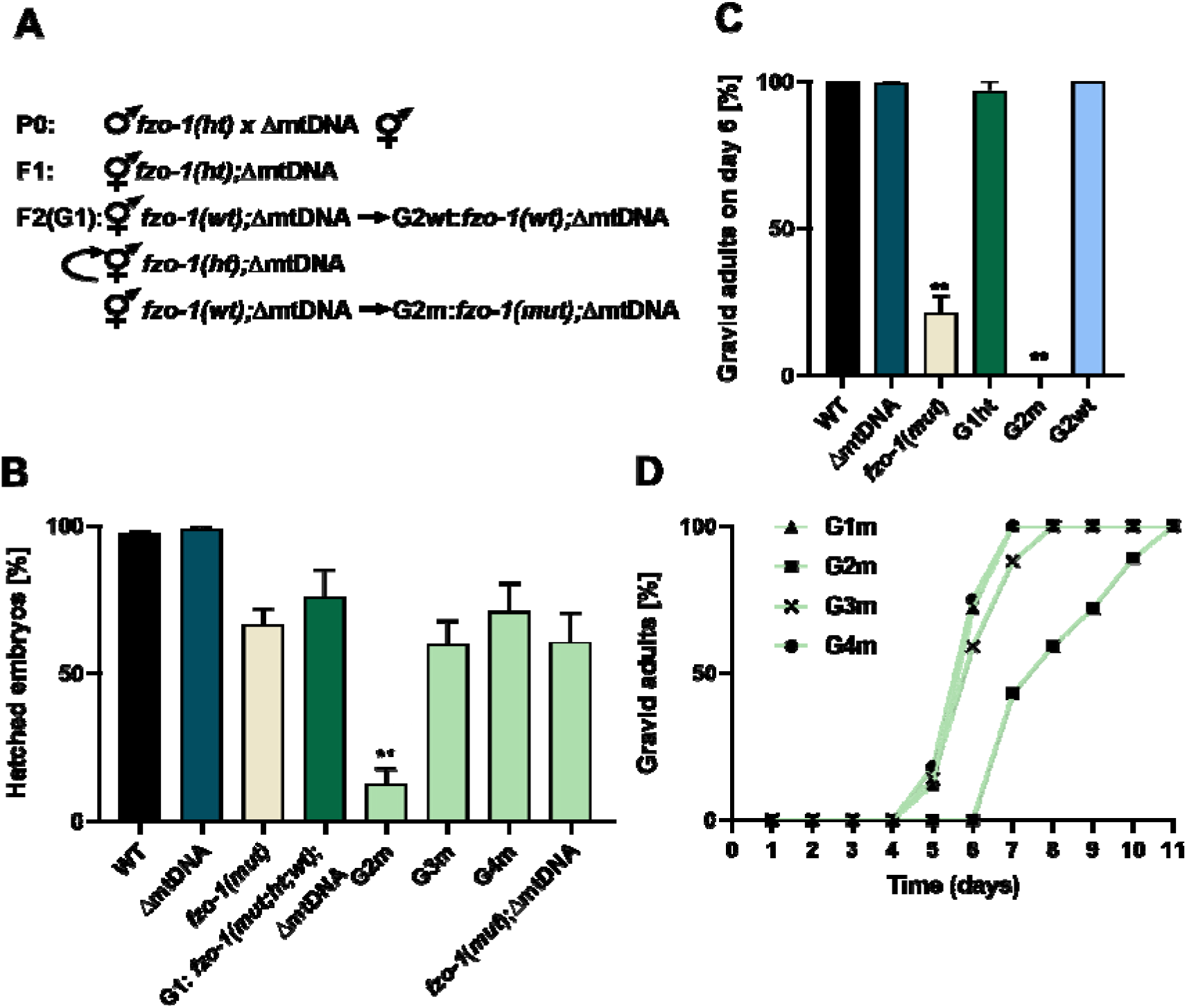
ΔmtDNA and *fzo-1(mut)* cannot co-exist. (A) Schematic representation of experimental setup. Heteroplasmic hermaphrodites (ΔmtDNA) were crossed with *fzo-1(mut)* males. Cross progeny F1 were allowed to self-propagate and single F2 (generation 1; G1) animals were isolated, allowed to lay eggs and their genotypes were determined. We then monitored heteroplasmic mutant or wild type *fzo-1* progeny over several generations (G2m-G4m and G2wt-G4wt, respectively). Heterozygous progeny was maintained to generate generation G1 without additional crosses. (B) The percent of hatched embryos of parental strains: N2(WT), ΔmtDNA and *fzo-1(mut)*, of *fzo-1(mut)* mutant cross progeny across generations (G1m-G4m) and of the stable cross line (>20 generations). P values were calculated using the Wilcoxon Mann-Whitney rank sum test by comparison with *fzo-1(mut)* animals. (**) denotes P<0.01. (C) The percent of gravid adults six days after egg laying of parental strains: N2(WT), ΔmtDNA and *fzo-1(mut)*, and of mutant cross progeny across generations (G1m-G4m). P values were calculated using the Wilcoxon Mann-Whitney rank sum test by comparison with *fzo-1(mut)* animals. (**) denotes P<0.01. (D) The percent of gravid adults of mutant cross progeny across generations (G1m-G4m) at the indicated times after egg laying.

This was in contrast to ~75% of the G1 animals that reached adulthood (Fig. 1C). These findings demonstrate that the interaction between the heteroplasmic ΔmtDNA and the nuclear DNA-encoded *fzo-1* mutant led to a severe reduction in fitness.

### The adverse effects of interaction between ΔmtDNA and *fzo-1(mut)* are reversed across generations

To better characterize the phenotypic impact of the interactions between ΔmtDNA and *fzo-1(mut)*, we monitored the progeny of the self-crossed *fzo-1(ht);*ΔmtDNA worms. Specifically, we measured the duration of the larva-to-adulthood period during the course of development in the G1m-G4m generations (Fig. 1D). While ~75% of the G1m reached adulthood after six days, the development of G2m animals was severely delayed, with 75% of the animals reaching adulthood only after nine days. To our surprise, the G3m animals showed significant improvement, with ~60% of this population reaching adulthood after six days. Moreover, G4m animals showed a full reversal of ΔmtDNA-associated adverse effects (Fig. 1D). We noted a similar pattern across generations when hatching was considered. In contrast to the 13.5% hatching observed among G2m embryos, 60±8% of the G3m embryos hatched. Strikingly, the hatching percentage of the G4m generation was indistinguishable from that of G1m animals and remained stable over subsequent generations (Fig. 1B). Finally, no phenotypic changes were observed across G1-G4 animals while tracing the developmental pace and hatching percentage of *fzo-1(wt)*;ΔmtDNA, G1wt-G4wt (Fig. S1). Taken together, our findings demonstrate a full reversal of the adverse effects of the interaction between the ΔmtDNA and the nuclear DNA-encoded mutant *fzo-1* gene.

### Mitofusin mutant leads to selection against ΔmtDNA heteroplasmy in *C. elegans*

We next asked how the deleterious interactions between *fzo-1(mut)* and ΔmtDNA were abrogated. We hypothesized that if the ΔmtDNA is not tolerated in the background of *fzo-1(mut)*, then selection against the ΔmtDNA should occur. To test this prediction, we quantified the levels of ΔmtDNA by quantitative PCR (qPCR) across the G1m-G4m generations (at the adult stage) in both the *fzo-1(mut);*ΔmtDNA and the *fzo-1(wt);*ΔmtDNA strains. We found that ΔmtDNA levels declined two-fold in the G1m *fzo-1(mut)* animals, as compared to *fzo-1* heterozygotes. This trend was enhanced, reaching a 10-fold reduction of ΔmtDNA levels in G2m animals. Values reached below detection levels in most G3m (N=19/22) and G4m (N=20/21) animals (Fig. 2A and S2A). In contrast, ΔmtDNA levels did not significantly change across the G1wt-G4wt generations of *fzo-1(wt);*ΔmtDNA animals (Fig. 2B and S2B).

**Figure 2:**
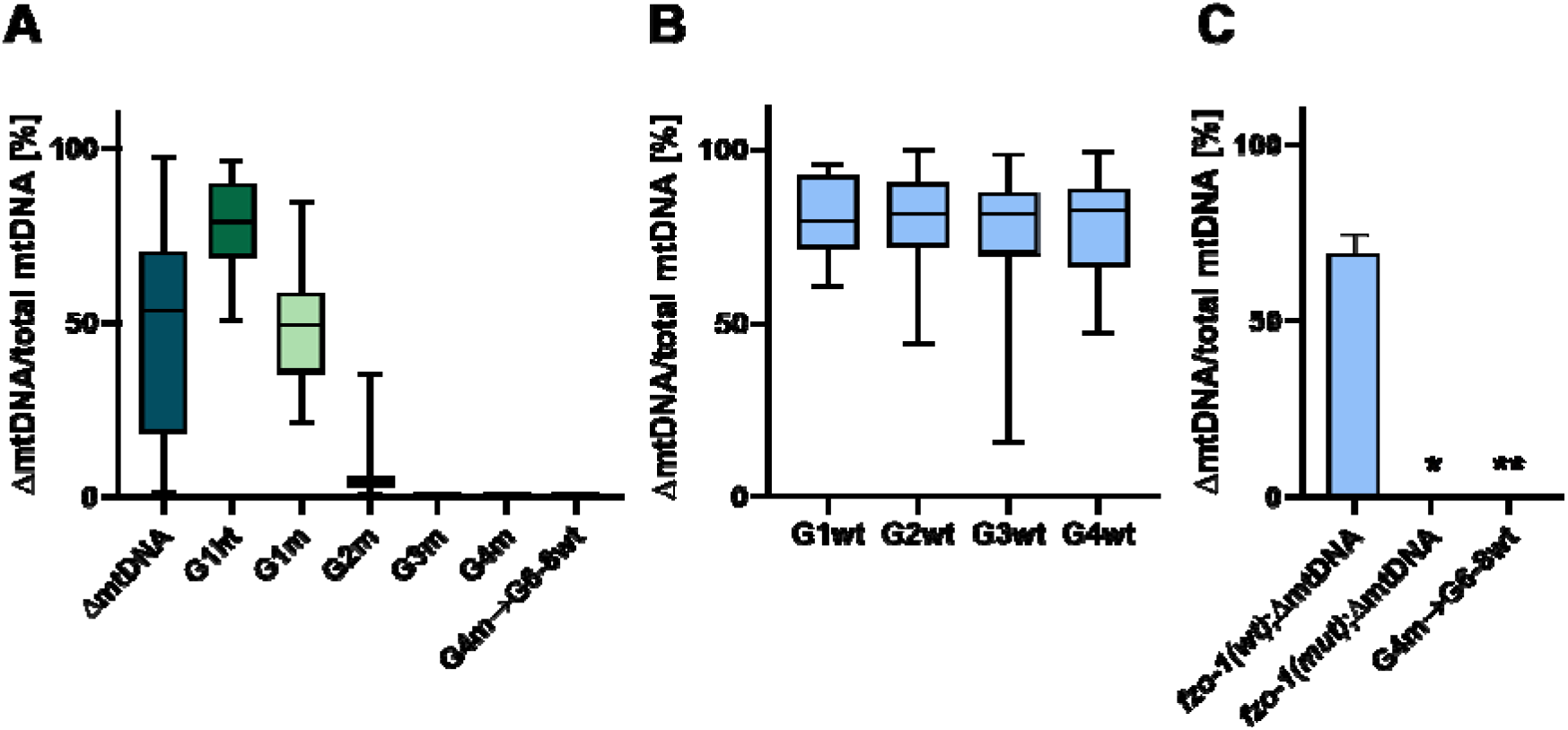
ΔmtDNA levels are selectively eliminated in *fzo-1(mut)*;ΔmtDNA animals across generations. (A) Box plot showing the percent of ΔmtDNA determined in individual animals (n≥20) of the parental heteroplasmic strain ΔmtDNA, the heteroplasmic *fzo-1(mut)* mutant cross-progeny strains (F1(ht), G1m-G4m) and the progeny of G4m animals crossed with *fzo-1(wt)*, (G4m->G6-8wt). (B) Box plot showing the percent of ΔmtDNA determined in individual animals (n≥15) of the heteroplasmic *fzo-1(wt)* cross progeny strains (G1wt-G4wt). (C) The percent of ΔmtDNA determined in a population of animals of the stable cross lines (>20 generations). P values were calculated using the Wilcoxon Mann-Whitney rank sum test by comparison with *fzo-1(wt);*ΔmtDNA animals. (*) denotes P<0.05 and (**) denotes P<0.01.

These results suggest that the ΔmtDNA was completely lost during the G1m-G4m generations. To test this hypothesis, we crossed G4m hermaphrodites with wild type males to isolate *fzo-1(wt)* progeny (G6). Since traces of ΔmtDNA were detected neither in G6 animals nor in subsequent generations (Fig. 2A and 2C), we concluded that disrupting *fzo-1* function indeed resulted in a complete and specific loss of the deleterious heteroplasmic ΔmtDNA. These results provide a proof of concept that mitochondrial fusion is critical for regulating the transmission of the *uaDf5* ΔmtDNA heteroplasmy across generation.

### Heteroplasmic truncations are differentially selected against based on size and mtDNA position in *C. elegans* mitofusin mutant

We next asked whether the deleterious interactions between *fzo-1(mut)* and ΔmtDNA depends on the size or genomic location of the mtDNA deletion. To achieve this goal, we characterized two additional mtDNA deletions obtained from the Million Mutation Project strain collection [27]. Specifically, these deletions comprise two new stable heteroplasmic *C. elegans* strains: *bguDf1* (derived from strain VC40128), harboring a mixture of intact mtDNA along with mtDNA molecules lacking ~1 kb (1kbΔmtDNA) encompassing two essential mtDNA genes (i.e., mt-ATP6 and mt-ND2); the second strain, *bguDf2* (derived from VC40469), harbors in addition to the intact mtDNA, a ~4.2 kb mtDNA deletion (4kbΔmtDNA) encompassing four different essential genes (i.e., mt-CO1, mt-CO2, mt-ND3 and mt-ND5). Notably, the levels of the 1kbΔmtDNA and 4kbΔmtDNA were stable over >100 generations (80% and 55%, respectively). At these heteroplasmy levels the animals displayed neither embryonic nor developmental phenotypes (Fig. S3A and S3B).

Heteroplasmic hermaphrodites of both strains were separately crossed with *fzo-1(mut)* heterozygote males, followed by F1 progeny self-cross (cross as in Fig. 1A). Similar to the approach taken with the *uaDf5* strain, we examined the distribution of the genotypes in the F2 heteroplasmic progeny. The genotypes distribution for the 1kbΔmtDNA did not deviate from the expected Mendelian ratios (23.5% homozygous *fzo-1(mut)*, 43.2% *fzo-1* heterozygotes (*ht*) and 33.3% *fzo-1(wt)* (p=0.14, Chi square test; Table S1). In contrast, these ratios strongly deviated from the expected Mendelian ratios for *fzo-1(mut)* animals harboring 4kbΔmtDNA (6.4% homozygous *fzo-1(mut)*, 59.3% *fzo-1* heterozygotes (*ht*) and 34.3% *fzo-1(wt)* (p=0.000076, Chi square test; Table S1). Hence, these results indicate that *fzo-1(mut)* differentially tolerates mtDNA deletions based on size and mtDNA position.

We then quantified the levels of ΔmtDNA in mutant versus wild type *fzo-1* progeny across four subsequent generations. We found that both truncated mtDNA molecules were undetectable after four generations, yet the decline rates diverged (Fig. 3A and 3B). Specifically, while the 1kbΔmtDNA was still detected in most of G3m animals (N=13/18), only a single G3m animal (N=1/21) still harbored the 4kbΔmtDNA molecule (Fig. 3C). It is worth noting that the levels of both types of ΔmtDNA did not significantly change across the G1wt-G4wt generations of the *fzo-1(wt);*ΔmtDNA animals (Fig. S3C and S3D).

**Figure 3:**
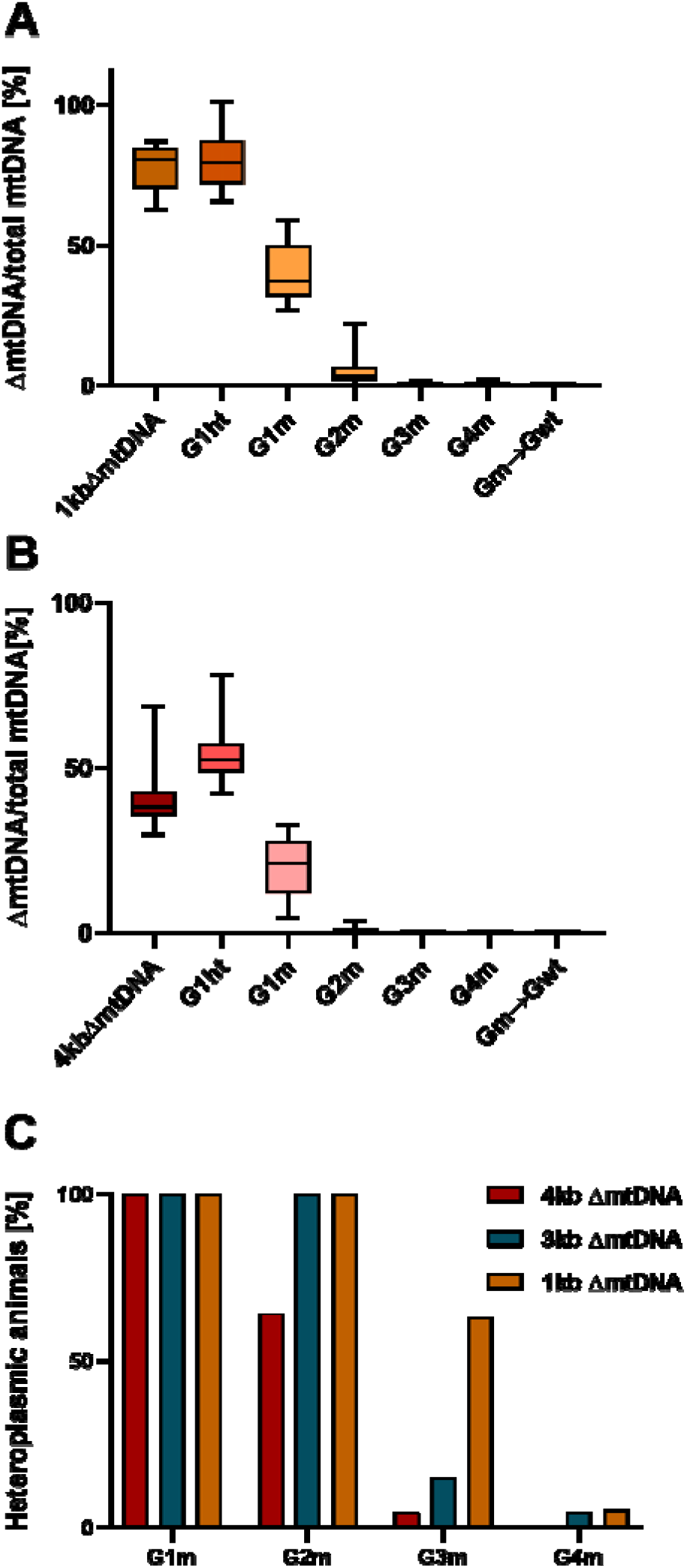
ΔmtDNA levels are differentially eliminated in *fzo-1(mut)*;ΔmtDNA animals based on size and mtDNA position. (A-B) Box plot showing the percentage of 1kbΔmtDNA (A) or 4kbΔmtDNA (B) ΔmtDNA, determined in individual animals (n≥14) of the parental heteroplasmic strain, the *fzo-1(mut)* mutant cross-progeny strains (F1(ht), G1m-G4m) and the progeny of G4m animals crossed with *fzo-1(wt)*, (G4m->G6-8wt). (C) The percentage of animals in each generation (G1m-G4m) carrying 1kbΔmtDNA (yellow), 3kbΔmtDNA (blue) or 4kbΔmtDNA (red; threshold was set at >0.01 AU).

To assess whether the truncated mtDNAs were completely lost, we crossed G4m hermaphrodites with wild type males and isolated *fzo-1(wt)* progeny (G6). qPCR analyses revealed no traces of the ΔmtDNA copies in subsequent generations (Fig. 3A and 3B). Thus, disrupting *fzo-1* function resulted in a complete and specific loss of a variety of heteroplasmic ΔmtDNAs. We interpret these results to mean that *fzo-1* function is sensitive to size and location of deleterious mtDNA heteroplasmy.

### ΔmtDNA molecules are selected against during *C. elegans* gametogenesis

The levels of an mtDNA heteroplasmic mutant were specifically reduced in the fly germline [20]. In *C. elegans*, mtDNA copy numbers start to increase significantly at the fourth larval stage (L4) is association with oocyte production [25, 28]. We, therefore, asked at which point during the *C. elegans* life cycle selection against ΔmtDNA occurred. Given that the relative levels of ΔmtDNA are maintained during development [25], we compared relative ΔmtDNA levels between embryos and adults in G2m animals. Our results indicate that relative ΔmtDNA levels were dramatically reduced (~5-fold) during development of G2m animals but not in G2wt animals (Fig. 4A and 4B). This observation suggests that ΔmtDNA is most likely selected against during worm development. Secondly, ΔmtDNA levels became undetectable in the resultant embryos of G2m animals (i.e., in G3m animals; Fig. 4A). Hence, it is possible that selection against ΔmtDNA molecules occurred during gametogenesis. In support of this claim, we noted that upon isolation of the gonads of G2m animals and then comparing ΔmtDNA levels in gonads to somatic tissues, we found a two-fold decrease in ΔmtDNA levels in the gonads. In contrast, no such difference was observed when comparing gonads and somatic tissues of *fzo-1(wt)*/ΔmtDNA animals (Fig. 4C and S4). Consistent with the *Drosophila* experiments [20, 29] and based on these observations, we suggest that selection against ΔmtDNA molecules during gametogenesis is evolutionarily conserved.

**Figure 4:**
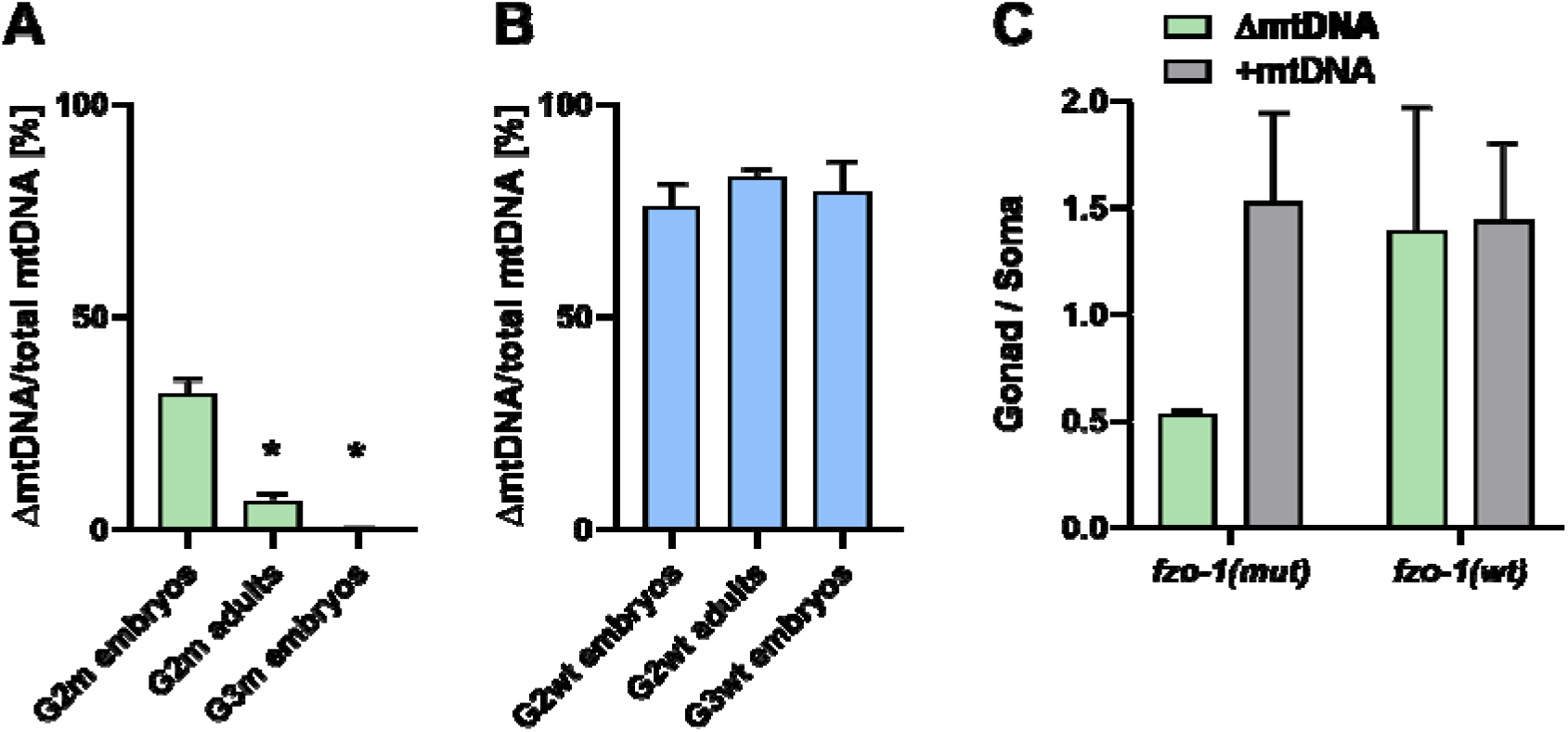
ΔmtDNA molecules are selectively eliminated during *C. elegans* gametogenesis. (A) The percent of ΔmtDNA in embryos and adults of the G2m-G3m generations. P values were calculated using the Wilcoxon Mann-Whitney rank sum test by comparison with *fzo-1(mut)* G2 embryos. (*) denotes P<0.05. (B) The percent of ΔmtDNA in embryos and adults of the G2wt-G3wt generations. P values were calculated using the Wilcoxon Mann-Whitney rank sum test by comparison with *fzo-1(wt)* G2 embryos. (C) The ratio of ΔmtDNA or +mtDNA levels in the gonad and soma of generation G2m adults. P values were calculated by comparison with *fzo-1(mut)* G2 embryos. (*) denotes P<0.05.

### PARKIN mutant aggravates the adverse effects of the ΔmtDNA-*fzo-1* interactions

Since Parkin mediates the turnover of mitofusins, and hence impacts their activity [30], we asked what would be the impact of the *pdr-1;fzo1* double mutant on the inheritance of the ΔmtDNA. To address this question, we first crossed ΔmtDNA heteroplasmic hermaphrodites with *pdr-1(gk448)* (here named *pdr-1(mut)*) males and established a stable *pdr-1(mut)* strain harboring heteroplasmy. In consistence with previous findings, the ΔmtDNA levels were elevated in this strain (75%) [13–15]. To establish a strain which is mutant in *pdr-1* and *fzo-1* in the context of ΔmtDNA heteroplasmy, we then crossed *pdr-1(mut)* heteroplasmic hermaphrodites with *pdr-1;fzo-1* heterozygous males, let the F1 progeny self-cross and isolated *pdr-1(mut);fzo-1(ht)* hermaphrodites harboring ΔmtDNA (Fig. 5A). This strain was allowed to propagate and the genotypic distribution of the heteroplasmic progeny was examined. While the genotype distribution of *fzo-1(mut)*; ΔmtDNA did not deviate from the expected Mendelian ratios (Table S1), the genotype distribution of *pdr-1;fzo-1;*ΔmtDNA was strongly affected, as follows: 9% homozygous *pdr-1(mut); fzo-1(mut)*, 55.8% *pdr-1(mut);fzo-1(ht)* and 35.2% *pdr-1(mut);fzo-1(wt)* (p=0.00055, Chi square test; Table S1). This suggests that the heteroplasmic ΔmtDNA cannot be tolerated in the background of *fzo-1;pdr-1* double mutant, as reflected in further reduction in fitness.

**Figure 5:**
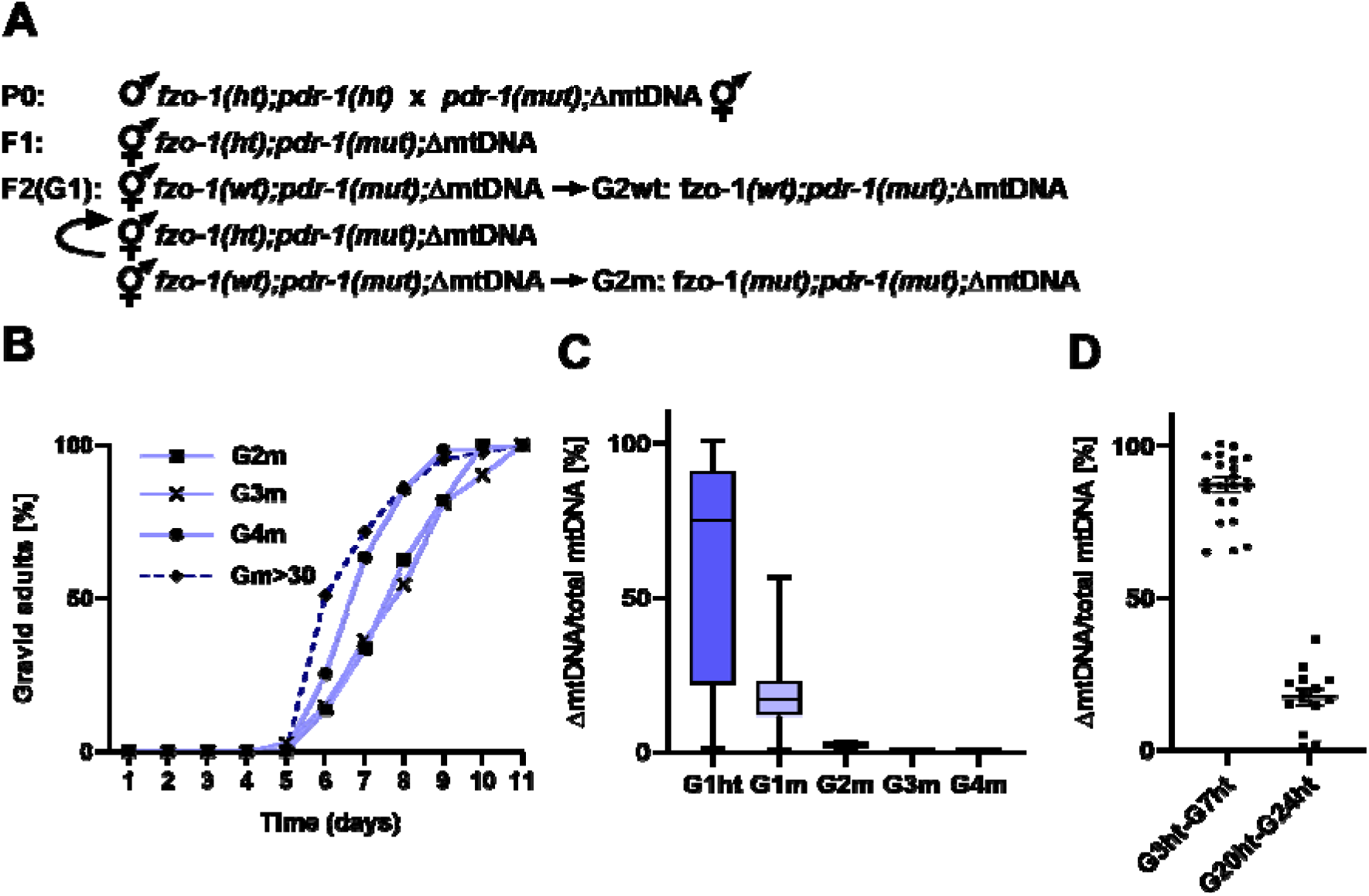
Mutant Parkin aggravates the adverse effects of the ΔmtDNA;*fZo-1*. (A) Schematic representation of experimental setup. Heteroplasmic *pdr-1(mut)* hermaphrodites (*pdr-1(mut);*ΔmtDNA) were each crossed with *fzo-1(ht);pdr-1(ht)* males. F1 progeny of such cross were allowed to self-propagate and single F2 animals (generation 1; G1) were isolated. These animals were allowed to lay eggs and their genotypes were determined. We then monitored the progeny of Δ*mtDNA;pdr-1(mut);fzo-1(mut)* or Δ*mtDNA;pdr-1(mut);fzo-1(wt)* animals over several generations (G2m-G4m and G2wt-G4wt, respectively). Δ*mtDNA;pdr-1(mut);fzo-1(ht)* progeny was maintained to generate generation G1 without additional crosses. (B) The percent of gravid adults of mutant cross progeny were monitored across generations (G2m-G4m) and of the stable cross line (>30 generations, Gm>30) at the indicated times after egg laying (C). Box plot showing the percent of ΔmtDNA was determined in individual animals (N≥14) of the Δ*mtDNA;pdr-1(mut);fzo-1(mut)* mutant cross-progeny strains (F1(ht), G1m-G4m). (D) Heteroplsamy levels of individual Δ*mtDNA;pdr-1(mut);fzo-1(ht)* animals sampled at generations 3-7 (Ght3-Ght7) and generations 20-24 (Ght20-Ght24).

We further monitored the development and fecundity of these animals. Phenotypic characterization revealed developmental delay in the *pdr-1(mut);fzo-1(mut)* G1m, similar to the parental strain (11/11 were adults by day 7); and most of the G1 m progeny (G2m eggs) hatched (85±8%). However, 66±14% of the G2m were developmentally arrested. Only 33±12% of the remaining animals reached adults by day 7, and their progeny production was severely reduced (laying 7 eggs or less over 20 hours). Moreover, while only 22±7.5% of the G3m animals were developmentally arrested, G3m development was similarly delayed (Fig. 5B and S5). However, during the G4m, significant recovery of animals’ development and egg hatching was noticed (Fig. 5B and S5). In contrast, heteroplasmic *pdr-1(mut);fzo-1(wt)* hatching and development was unaffected (~98% hatched and 100% adults by day 7; Fig. S5A and S5B). Thus, the adverse effects of the interaction between *fzo-1(mut)* and ΔmtDNA on fecundity and developmental timing were aggravated by *pdr-1(mut)*.

We next asked whether the selection against ΔmtDNA would be stronger in the background of *fzo-1(mut);pdr-1(mut)*. To directly examine this, we compared the levels of ΔmtDNA molecules in *fzo-1(mut);pdr-1(mut)* (Fig. 5C) to *fzo-1(mut)* (Fig. 2A) across G1m-G4m generations. We found that ΔmtDNA levels declined more sharply in *fzo-1(mut);pdr-1(mut)* as compared to *fzo-1(mut)*. Specifically, *fzo-1(mut);pdr-1(mut)* had 3-fold and 5-fold less ΔmtDNA in G1m and G2m, respectively. By G3m the heteroplasmy in all *fzo-1(mut);pdr-1(mut)* animals (N=16) was below detection levels (Fig. 5C). In contrast, ΔmtDNA levels did not significantly change across generations of *fzo-1(wt);pdr-1(mut)* ΔmtDNA animals (Fig. S5D). Taken together, the concomitant disruption of fusion and mitophagy strongly selected against ΔmtDNA molecules. In support of this interpretation, we noticed that even in the *fzo-1(ht);pdr-1(mut)* animals, where only one genomic copy of *fzo-1* remained functional, ΔmtDNA levels declined and in some individuals were lost across ~25 generations (Fig. 5D).

### MFN2 likely modulates heteroplasmic patterns in CMT2A patients

To assess the potential impact of defective MFN2 on the dynamics of mitochondrial heteroplasmy in humans, we employed massive parallel sequencing (MPS) of the entire mtDNA in CMT2A patients and healthy maternal relatives in three independent pedigrees (Fig. S6). High coverage per mtDNA nucleotide (>1000X) was attained at a mean of 16560 mtDNA positions, thus allowing both assignment of samples to specific mtDNA genetic backgrounds (haplogroups) and the detection of heteroplasmy levels >1% (Table S2).

In pedigree 1, a Caucasian-Cherokee American Indian family in which a Leu146Phe MFN2 mutation [31] has been segregating, five heteroplasmic mutations were identified, two of which (mtDNA positions 310 and 16172) are shared between the female patient (sample V17) and her healthy siblings (samples V13 and V14; Table S3). Interestingly, although the C-to-G transition at position 16172 was highly prevalent (~95%) in the healthy maternal siblings, it became rare in the patient (~5%). Notably, position 16172 maps within the mtDNA transcription termination site (TAS) [32], thus suggesting potential functionality.

In pedigree 2 (Caucasian family 1706, with the L76P MFN2 mutation [33]), a heteroplasmic transition at mtDNA position 13830 was identified (a synonymous mutation in the ND5 gene), which differentially segregated among patients and healthy maternal relatives. Nevertheless, the ratio between the two segregating alleles at this position was the opposite of what was seen between the proband (individual 106) and his maternal uncle, who was also a patient (individual 1007). This finding is consistent with random segregation of this heteroplasmic mutation. Finally, analysis of the entire mtDNA sequence in patients and healthy maternal relatives in pedigree 3 (an Arab-Israeli family presenting the Q386P MFN2 mutation [34]) did not reveal any trustworthy heteroplasmic mutations. Thus, we found that only in the case where a heteroplasmic mutation was potentially functional was its level significantly declined exclusively in the CMT2A patient. We speculate that selective heteroplasmy removal could have occurred, consistent with our *C. elegans* observations.

## Discussion

Here, by manipulating the fusion machinery in *C. elegans*, we demonstrated that mitofusin (f*zo-1*) is a key modulator of mtDNA heteroplasmy, and show its impact on transmission of heteroplasmic deletions across generations in living animals. Firstly, we discovered that a mitofusin mutant led to complete loss of three different mtDNA deletions, separately. These findings provide experimental support for the hypothesis that functional complementation among mitochondria in the intracellular network likely enables the survival and prevalence of deleterious heteroplasmic mtDNA mutations in the population [23, 24]. Secondly, we found that the mitofusin mutant was differentially sensitive to the size and location of the different deletions in the mitochondrial genome. Third, analysis of *fzo-1;pdr-1* double mutants selected against the survival of animals with heteroplasmic mtDNA deletion, and accelerated the loss of truncated mtDNA molecules across generations. Taken together, our results demonstrate the importance of cross talk between mitochondrial fusion and the mitochondrial quality control machineries in protecting living animals from the adverse impact of inheriting deleterious mtDNA molecules.

Our discovery that heteroplasmic deletions are lost across generations when the fusion machinery is disrupted is consistent with previous findings of selective forces in the human germline [4, 5]. Moreover, in *Drosophila*, germline selection of heteroplasmic mutations was directly observed in response to compromised fusion and quality control machineries [20, 29]. Our analyses build upon these observations, and followed the dynamics of heteroplasmy in multiple subsequent worm generations, which enabled identifying complete loss of the heteroplasmy. Therefore, heteroplasmy of deleterious mutations cannot be tolerated unless in the presence of functional compensatory mechanism inherent to the mitochondrial network, as well as the quality control machinery.

Our analysis of three different mtDNA deletions in *C. elegans* revealed differences in the pace of their loss, and levels of phenotypic severity, when grown in the presence of a mitofusin mutant (Fig. 6). Indeed, these three deletions encompass different sets of mtDNA genes, and they differ in size, with the latter seemingly playing a more pronounced role in the differences between the survival rates of the worms. This not only underlines the importance of intra-cellular mitochondrial network in protecting against the deleterious impact of heteroplasmic mutations, but also suggests that the fusion machinery is sensitive to differential severity of the phenotypic impact of heteroplasmic mutations. This finding is in line with differences in the penetrance of disease-causing mutations, which ranges between 60-80%, depending on the symptoms [9]. Nevertheless, the question about the functional importance of certain mtDNA regions versus others remain open. This calls for a screen of mtDNA mutants that will systematically enable assessing sensitivity of the mitochondrial quality control and fusion machineries in differentiating the phenotypic impact of a variety of mutations, locations and sizes.

**Figure 6:**
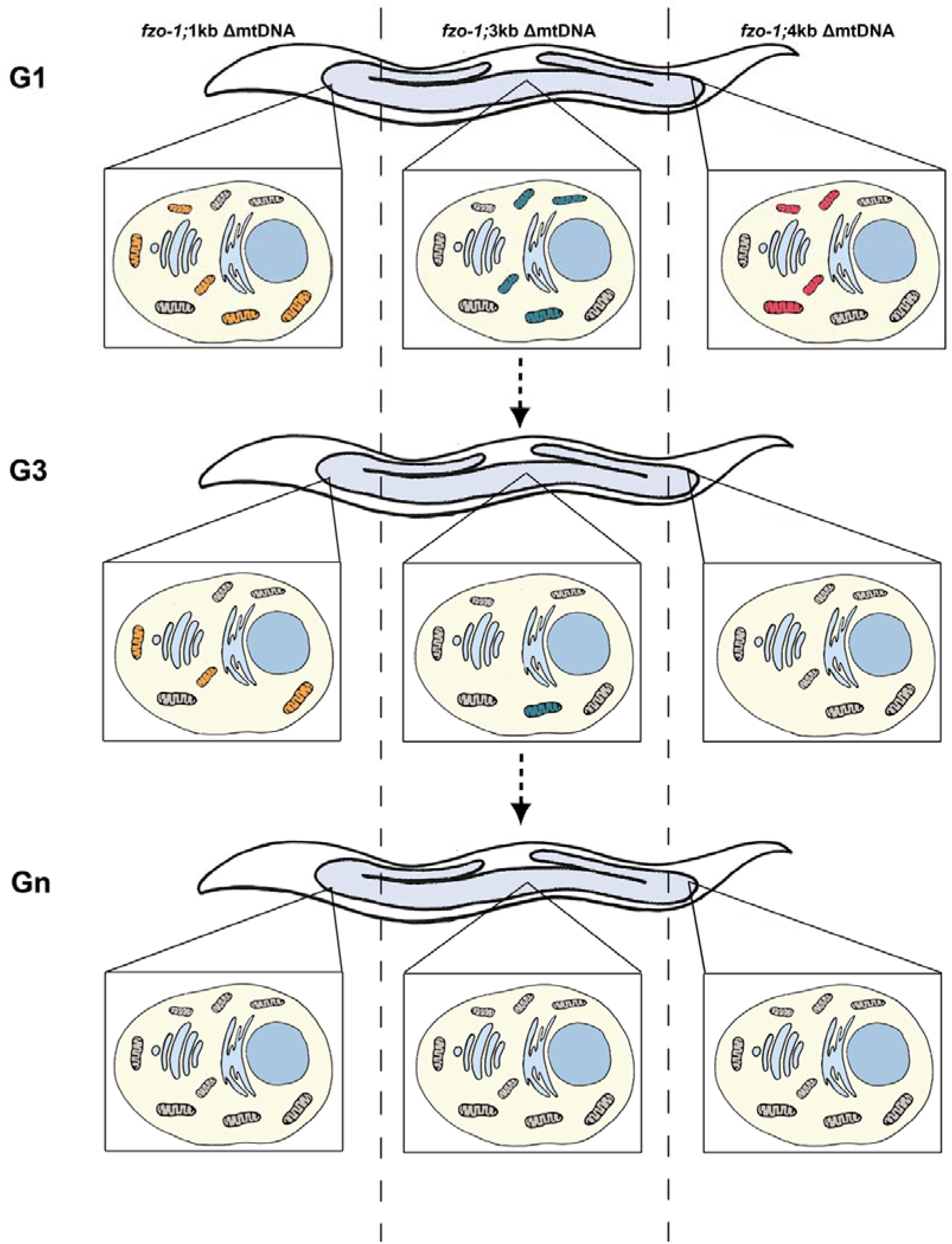
A model depicting the separate selective removal of three ΔmtDNA across generation in the *C. elegans*. Schematic representation of ΔmtDNA levels in three heteroplsamy strains, carrying intact mtDNA (black) and 1kbΔmtDNA (yellow), 3kbΔmtDNA (blue) or 4kbΔmtDNA (red) across generation. Differential sensitivity of the fusion machinery to the different heteroplasmic deletions is illustrated.

It was previously found that *fzo-1* mutation impacted the developmental pace of the worms [35–37]. Nevertheless, we found that the introduction of heteroplasmic mtDNA deletions strongly aggravated these phenotypes, leading to a sharp decline in survival and fecundity. This was manifested by reduced egg hatching, and delayed or even arrested larval development. This strong selective pressure positively correlated with the decline in heteroplasmic deletions’ levels across generation; the fact that the worms’ phenotype was ameliorated within 3-4 generations, upon loss of ΔmtDNA molecules suggests that the role of mitochondrial fusion in tolerating ΔmtDNA is critical for the fitness of the organism. Finally, we found that disrupting mitophagy in addition to mitochondrial fusion (*fzo-1;pdr1* double mutant) in the presence of ΔmtDNA sharply increased lethality already in G1 animals (reflected by the skew from Mendelian ratios). Therefore, we argue that the interaction between mitochondrial fusion and quality control machineries is not only critical to cope with ΔmtDNA within the cell, but is essential for the organism fecundity, development, and survival across generations.

If mitochondrial fusion is indeed important for modulating the inheritance of mtDNA heteroplasmy, one could anticipate that dysfunctional mitochondrial fusion, such as in the case of Charcot Marie Tooth type 2A (CMT2A) patients, will affect patterns of heteroplasmy. Our analysis of three CMT2A pedigrees lend first clues that this might be the case. Whereas deep mtDNA sequencing in two of the pedigrees did not reveal any potentially functional mtDNA mutations, we found that the levels of a potentially functional mtDNA mutation in a patient was notably lower than her healthy maternal relatives. We note that these results are in line with the observations in worms, supporting our working hypothesis that the fusion machinery modulates the levels of deleterious mtDNA heteroplasmy across generations to allow tolerance and survival. Future collection of a larger number of CMT2A pedigrees is required to draw clearer conclusion.

The complete loss of ΔmtDNA across *C. elegans* generations suggests that the fusion machinery represents an attractive candidate target for future treatment of mitochondrial disorders. For example, the activity of these two machineries decline during the aging of the individual [38], and the levels/repertoire of mtDNA heteroplasmic deletions increase in tissues from aged individuals [39]. It would, therefore, be of great interest to assess the importance of such three-way interaction (i.e., mitochondrial fusion-mitochondrial quality control and patterns of heteroplasmy) to age associated diseases.

## Methods

### Nematodes and growth conditions

A list of strains used in this work and name abbreviations is found in Table S4. All strains were outcrosses to our N2 stock at least four times. Nematodes were grown on Nematode Growth Medium (NGM) plates seeded with the *Escherichia coli* OP50-1 strain at 15°C.

### Crosses

Mutant *fzo-1(tm1133)* animals (strain CU5991) are very poor in mating and, therefore, were first crossed with males expressing a yellow fluorescent protein marker (*unc-54p::YFP*). Heteroplasmic hermaphrodites (*uaDf5/+, bguDf1/+* or *bguDf2/+*) were then crossed with *fzo-1(tm1133);unc-54p::YFP* heterozygote males to ensure maternal inheritance of mtDNA deletions and to establish independent heteroplasmic lines carrying the wild type or *fzo-1(tm1133)* mutation. Mutant *pdr-1(gk448)* animals (strain VC1024) were first crossed with heteroplasmic hermaphrodites (*uaDf5/+*) and with *fzo-1(tm1133)* to establish double mutant strains. *fzo-1(tm1133);pdr-1(gk448)* double mutants were crossed with *unc-54p::YFP;* then, the F1 heterozygote males, *fzo-1(tm1133);pdr-1(gk448);unc-54p::YFP, were* crossed with heteroplasmic hermaphrodites, ΔmtDNA *pdr-1(gk448)* to establish independent heteroplasmic lines carrying the *pdr-1(gk448)* mutation and either the wild type or *fzo-1(tm1133)* mutation. F2 progeny of all crosses were screened for *fzo-1(tm1133)* and/or *pdr-1(gk448)* mutations using a single worm PCR Phire Animal Tissue Direct PCR Kit (Thermo Scientific). The resultant amplification products were visualized by gel electrophoresis. List of PCR primers is found is Table S5. Heteroplasmic (*uaDf5/+, bguDf1/+* or *bguDf2/+*) animals that were heterozygotes for *fzo-1(tm1133)* were maintained to easily produce G1 animals.

### Embryo hatching

Gravid animals were moved to fresh plate for 2-12 hours and then removed from the plates. Embryos were allowed to develop and hatching was examined after 48 hours. Experiments were repeated at least four times, and >100 embryos were scored per experimental condition. P values were calculated using the Wilcoxon Mann-Whitney rank sum test to compare two independent populations.

### Developmental timing

Single embryos were placed on fresh plates and allowed to grow at 15°C. The animals’ developmental stage was examined every day and the number of animals reaching reproductive adulthood on each day was recorded. Developmentally arrested animals that did not reach adulthood in over 10 days were excluded. Experiments were repeated independently at least three times, and >30 animals were scored per experimental condition. P values were calculated using the Wilcoxon Mann-Whitney rank sum test to compare two independent populations.

### DNA purification and extraction

Total DNA was extracted using a QuickExtract kit (Lucigen). Unless otherwise indicated, DNA was extracted from a single worm (n≥14). When populations were examined, ~5 animals were collected. For embryos, DNA was extracted from ~30 embryos. For gonad-soma analysis, gonads were dissected from ~5 animals per biological repeat. DNA was extracted separately from the gonads and soma.

### Assessment of mtDNA copy numbers

mtDNA levels were measured by qPCR performed on a C1000 Thermal Cycler (Bio-Rad) with KAPA SYBRFAST qPCR Master Mix (KAPA Biosystems). Analysis of the results was performed using CFX Manager software (Bio-Rad). To quantify the different mtDNA molecules, three set of primers were used for truncated, intact and total mtDNA molecules for each of the three deletions examined (Table S5). The average C_T_ (threshold cycle) of triplicate values obtained for these mtDNA molecules was normalized to a nuclear DNA marker using the 2^-ΔΔC^_T_ method [40]. At least four independent experiments were used to determine the normalized C_T_ values of each strain or generation. Truncated/total ratio was defined as the ratio of the normalized C_T_ values of truncated to total mtDNA for a given strain.

### Gonad dissection

G2 wild type or mutant animals were placed in a drop of ultra-pure water on a coverslip slide and a 25-gauge needle was used to remove the gonads from the body of the animals. Gonads or the remaining carcasses were then transferred to DNA extraction buffer. 5-10 worms were used for each biological repeat. Experiments were repeated independently at least four times. P values were calculated using the Wilcoxon Mann-Whitney rank sum test to compare two independent populations.

### Human DNA samples

Patients diagnosed with Charcot-Marie-Tooth type 2a (CMT2A) and healthy maternal relatives from three unrelated pedigrees were considered in the current study. Total blood DNA was extracted from available patients and healthy individuals of all three sibs. Pedigree 1, of English-native American admixed origin, harbors patients heterozygous for the dominant L146F MFN2 mutation [31]. Pedigree 2, of North American Caucasian origin, harbors patients previously identified as heterozygous for the L76P MFN2 dominant mutation [33]. In pedigree 3 of Israeli Arabian origin, the patient was identified as a compound heterozygote for A1157C and T1158G mRNA nucleotide position mutations, both leading to the recessive Q386P mutation [34]. In Fig. S6, arrows point to the tested individuals in each pedigree. The samples were collected according to the accepted ethical regulations following the institutional IRBs.

### Massive parallel sequencing of the entire human mtDNA and bioinformatics analysis

The entire mtDNA of available patients and controls (Fig. S6) was amplified in three fragments using three sets of primers, as previously described [2]. For each of the analyzed samples, the amplified fragments were mixed in equimolar ratios and sent for library construction and sequencing utilizing the Illumina MiSeq platform (Technion Genome Center, Israel). Paired-end reads were trimmed using TrimGalore [41] (version 0.4.5) and then mapped to the mtDNA reference sequence (rCRS) using BWA mem (version 0.7.16) with default parameters [42]. Alignment files (SAM format) were compressed to their binary form (BAM format) using SAMtools (version 1.3) with default parameters [view –hb] and sorted using the [sort] function [43]. True heteroplasmic mutations were annotated using an in-house script that followed the logic of the previously established MitoBamAnnotator [44]. In brief, a pileup of the mtDNA-mapped reads was generated using the SAMtools [mpileup] function with the [-Q 30] parameter to only consider reads with a phred score higher than 30 as a measure of quality control. Next, the mapped bases were counted and the most frequent nucleotide in each mtDNA position was considered the major allele only if that nucleotide position had a minimal coverage of 10X. To avoid strand bias, the secondary mutation at a given nucleotide position was recorded only if it was represented by at least two reads per strand. mtDNA sequences were then haplotyped using HaploGrep2.0 [45].

### Sanger sequencing and cloning

Sanger sequencing was employed to verify the identified heteroplasmic mutations at position 16172 (pedigree 1) and position 13830 (pedigree 2). Primers used for this amplification reactions are listed in Table S6.

### NUMT exclusion

To control for possible contamination by nuclear mitochondrial DNA pseudogenes (NUMTs) in the MPS reads, mtDNA sequence contigs encompassing each of the identified mutations were BLAST screened as previously described [46]. In brief, the highest scored nuclear DNA BLAST hits encompassing each of the identified mutations were aligned against the mtDNA sequence reads of the relevant sample and the number of reads harboring the exact identified mutation was counted using IGV viewer [47]. Notably, as mtDNA is maternally inherited as a single locus, variants appear in linkage.

Hence, combinations of variants that were present in nuclear DNA BLAST hits but which were not identified in sequence reads of the relevant samples were not considered as corresponding to NUMTs.

## Acknowledgements

This study was funded by the Israel Science Foundation (ISF) grant 278/18 to ABZ and grant 372/17 to DM. L.M. was supported by the Ministry of Science and Technology, Yitzhak Navon PhD fellowship and Kreitman Biotech scholarship. D.B.Y. and T.C. were supported by the Kreitman Negev scholarships.

## Author Contributions

Conceptualization, A.B. and D.M.; Methodology, L.M., D.B. and I.V.; Investigation, L.M., D.B., D.K., T.N., M.K. and S.D.; Formal Analysis, D.B and T.C; Resources, C.J.K., J.M.V., Y.N. and S.Z; Writing, Review & Editing, A.B. and D.M.; Supervision, A.B. and D.M.; Funding Acquisition, A.B. and D.M.

## Data and availability

All supporting data is included in the manuscript and supporting information.

