## Supplementary Figures and Tables for "Mutant *C. elegans* mitofusin leads to selective removal of mtDNA heteroplasmic deletions at different rates across generations"

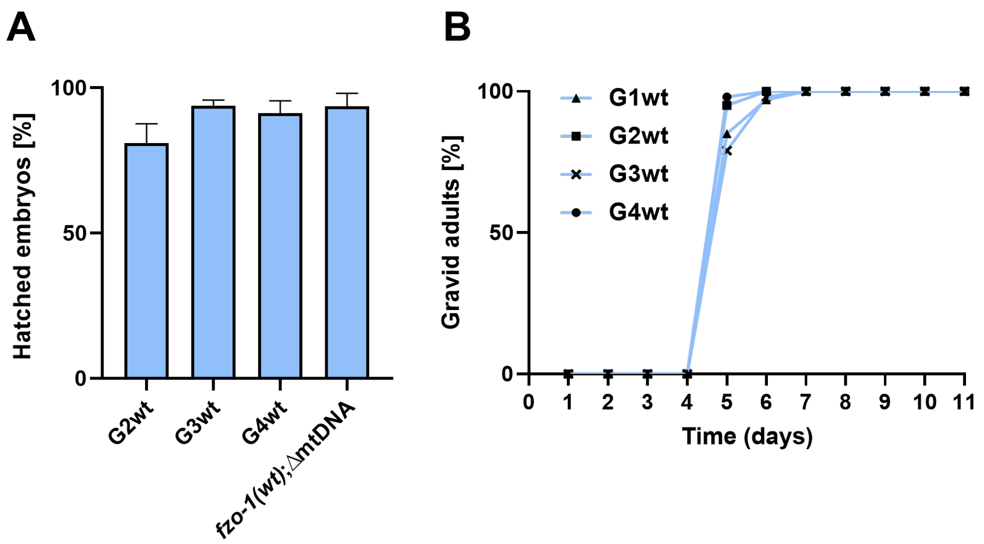


**Figure S1: Characterization of ΔmtDNA;*fzo-1(wt)* animals.** (A) The percent of hatched embryos of ΔmtDNA;*fzo-1(wt)* mutant cross progeny across generations (G1wt-G4wt). (B) The percent of gravid adults of ΔmtDNA;*fzo-1(wt)* cross progeny across generations (G1wt-G4wt) at the indicated times after egg laying. P values were calculated using the Wilcoxon Mann-Whitney rank sum test by comparison with G2wt animals.

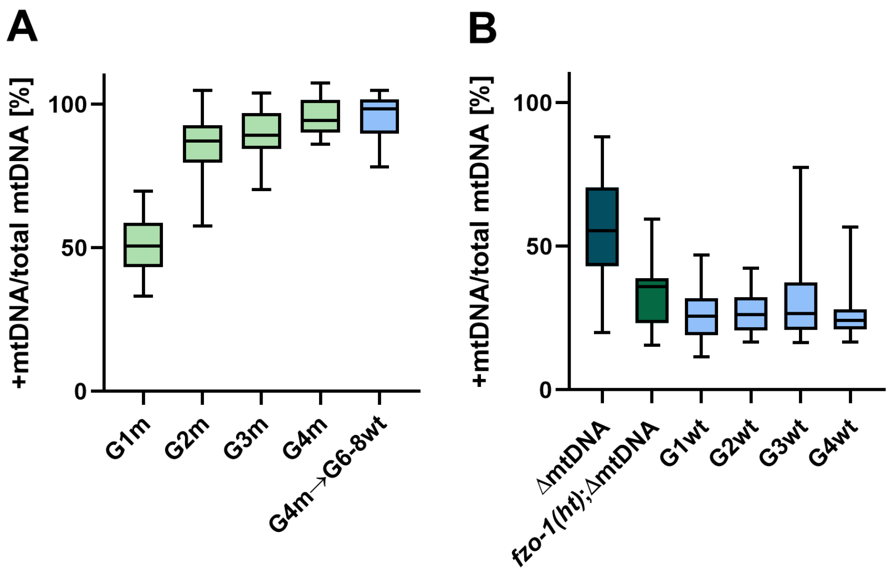


**Figure S2: +mtDNA levels across generations.** (A) Box plot showing the percent of +mtDNA determined in individual animals (n≥20) of the parental heteroplasmic strain ΔmtDNA, the *fzo-1(mut)* mutant cross-progeny strains (F1(ht), G1m-G4m) and the progeny of G4m animals crossed with *fzo-1(wt)*, (G4m->G6-8wt). (B) Box plot showing the percent of +mtDNA determined in individual animals (n≥15) of the *fzo-1(wt)* cross progeny strains (G1wt-G4wt).

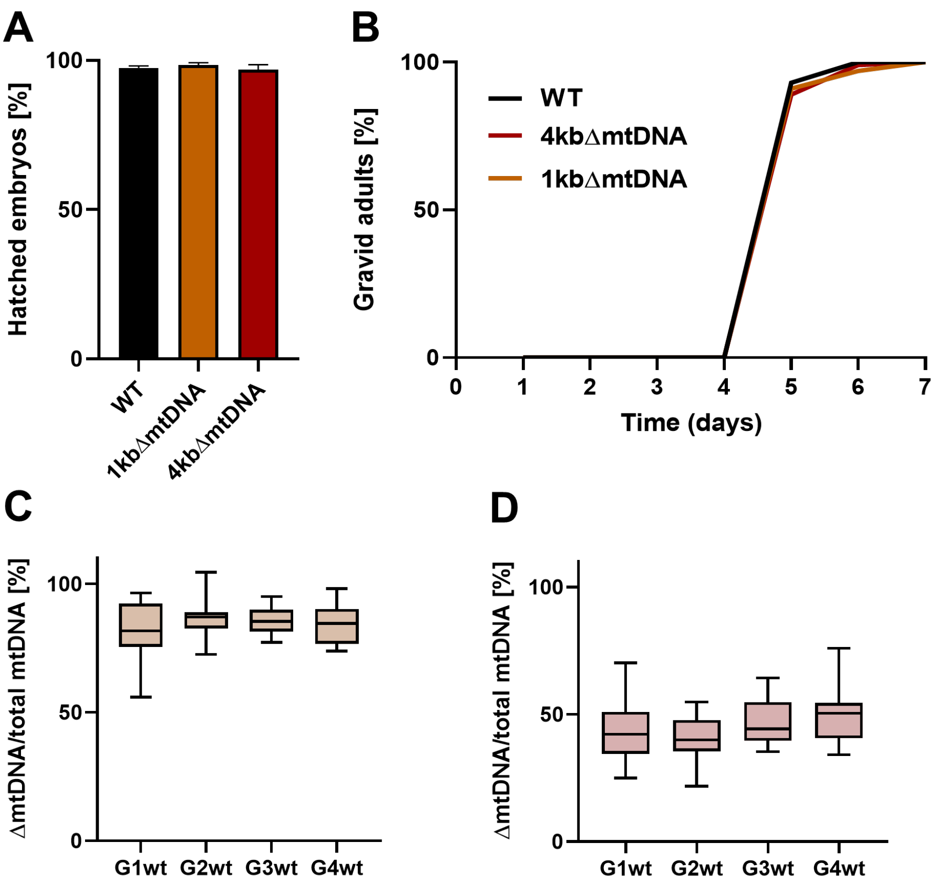


**Figure S3: Characterization of** the **1kbΔmtDNA and 4kbΔmtDNA strains**. (A) The percent of hatched embryos of N2 (WT), 1kbΔmtDNA and 4kbΔmtDNA. P values were calculated using the Wilcoxon Mann-Whitney rank sum test by comparison with N2 embryos. (B) The percent of gravid adults of N2 (WT), 1kbΔmtDNA and 4kbΔmtDNA animals at the indicated times after egg laying. (C-D) Box plot showing the percent of either 1kbΔmtDNA (C) or 4kbΔmtDNA (D) determined in individual animals (n≥15) of the heteroplasmic *fzo-1(wt)* cross progeny strains (G1wt-G4wt).

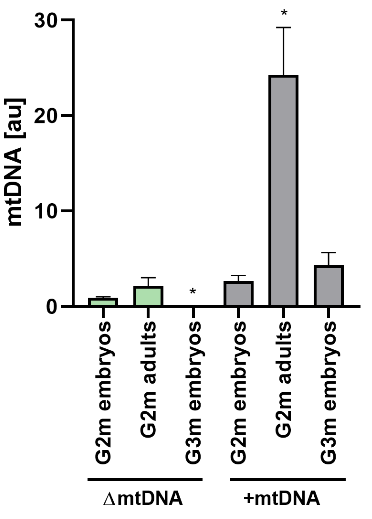


**Figure S4: ΔmtDNA levels do not significantly increase during development.** Levels of ΔmtDNA and +mtDNA in embryos and adults in the G2m-G3m generations. P values were calculated using the Wilcoxon Mann-Whitney rank sum test by comparison with G2m embryos. (*) denotes P<0.05.

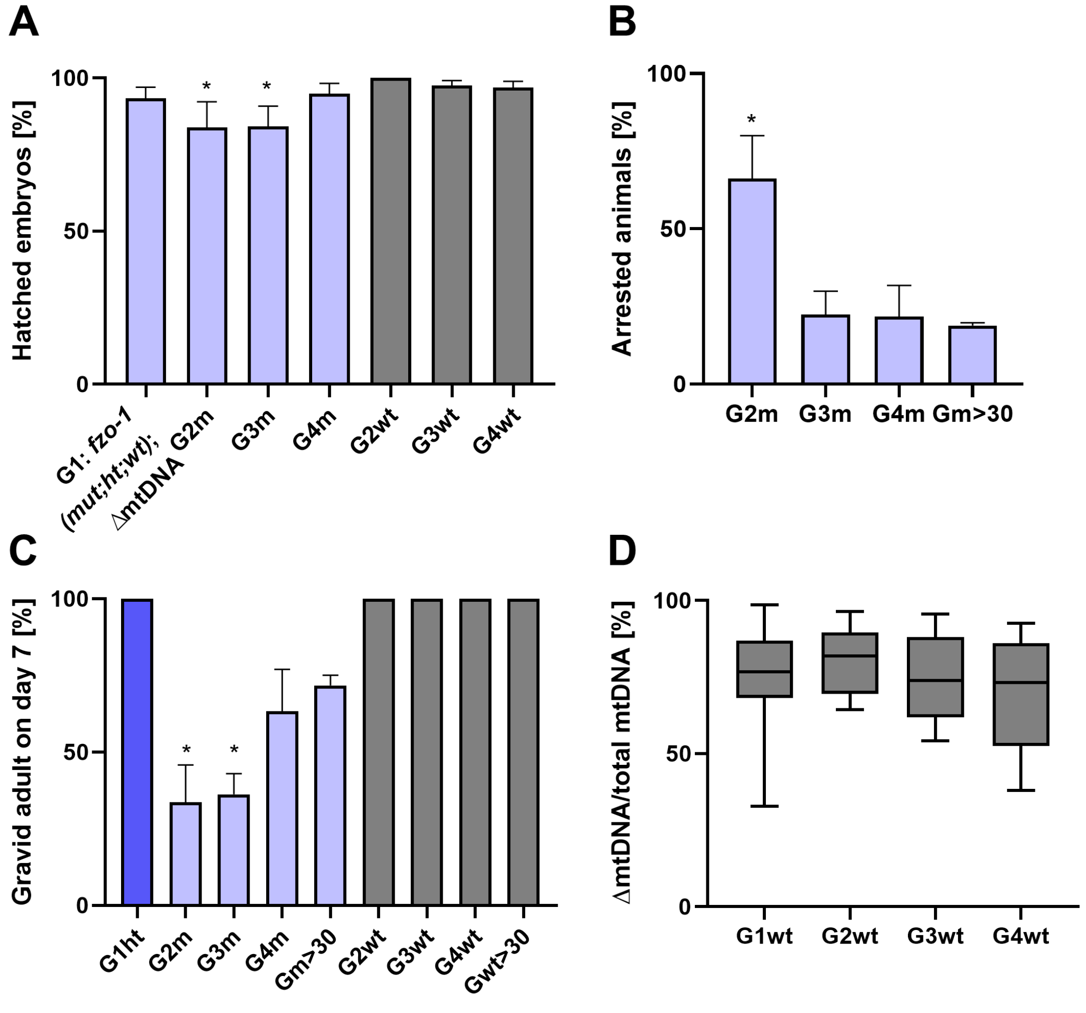


**Figure S5: Characterization of ΔmtDNA;*pdr-1(mut);fzo-1(wt)* animals.** (A) The percent of hatched embryos of heteroplasmic *pdr-1(mut);fzo-1(mut)* or *pdr-1(mut);fzo-1(wt)* cross progeny monitored across generations (G1ht, G2m-G4m and G2wt-G4wt). P values were calculated using the Wilcoxon Mann-Whitney rank sum test by comparison with G1ht embryos. (*) denotes P<0.05. (B) The percent of gravid adults of 7 days after egg laying of heteroplasmic *pdr-1(mut);fzo-1(mut)* or *pdr-1(mut);fzo-1(wt)* monitored across generations

(G1ht, G2m-G4m, G2wt-G4wt and stable lines (>30 generations), Gm>30, Gwt>30). P values were calculated using the Wilcoxon Mann-Whitney rank sum test by comparison with G1ht animals. (*) denotes P<0.05. (C) The percent of developmentally arrested animals of ΔmtDNA;*pdr-1(mut);fzo-1(mut)* mutant cross progeny across generations (G2m-G4m) and the stable line (>30 generations, Gm>30). P values were calculated using the Wilcoxon Mann-Whitney rank sum test by comparison with G4m animals. (*) denotes P<0.05. (D) Box plot showing the percent of ΔmtDNA determined in individual animals (N≥16) of the ΔmtDNA*;pdr-1(mut);fzo-1(wt)* cross-progeny strains (G1wt-G4wt).

**a**

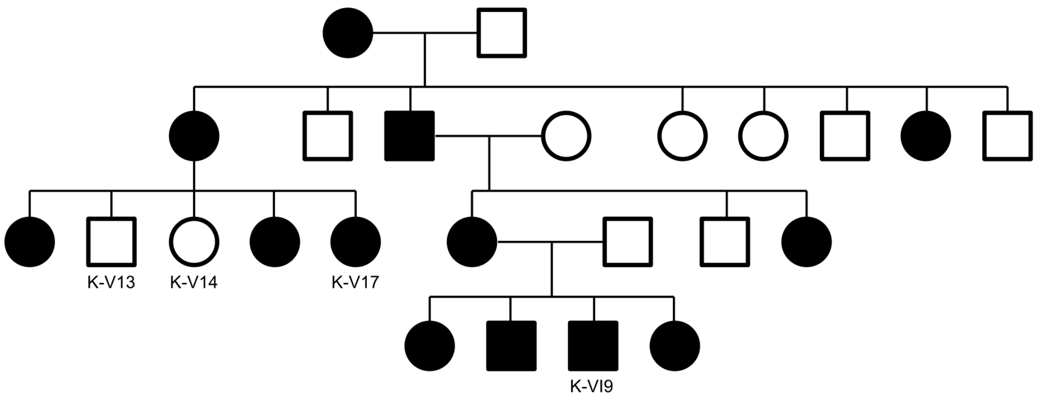


**b**

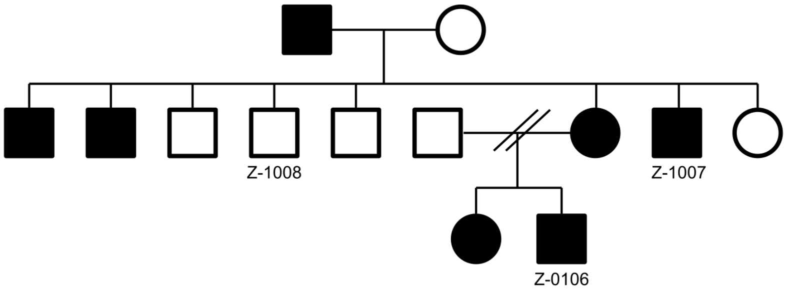


**c**

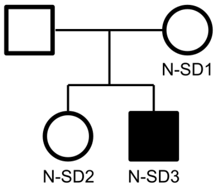

**Figure S6: Representative pedigrees of the analyzed CMT2A patients**. Three pedigrees depicting the inheritance of Charcot-Marie-Tooth disease (filled objects). The analyzed samples are indicated by name according to Table S3.

**Table S1: Genotypes distribution of F2 progeny in different heteroplasmic strains**

| Genotype | Progeny (#) | Progeny (%) | $\chi^2$ test (N) |
|---|---|---|---|
| *fzo-1(wt);ΔmtDNA* | 38 | 25.3 | 0.96 (150) |
| *fzo-1(ht);ΔmtDNA* | 73 | 48.7 | |
| *fzo-1(mut);ΔmtDNA* | 39 | 26 | |
| *fzo-1(wt);1kbΔmtDNA* | 61 | 33.3 | 0.14 (183) |
| *fzo-1(ht);1kbΔmtDNA* | 79 | 43.2 | |
| *fzo-1(mut);1kbΔmtDNA* | 43 | 23.5 | |
| *fzo-1(wt);4kbΔmtDNA* | 48 | 34.3 | 0.000076 (140) |
| *fzo-1(ht);4kbΔmtDNA* | 83 | 59.3 | |
| *fzo-1(mut);4kbΔmtDNA* | 9 | 6.4 | |
| *fzo-1(wt);pdr-1(mut);ΔmtDNA* | 70 | 35.2 | 0.00055 (199) |
| *fzo-1(ht);pdr-1(mut);ΔmtDNA* | 111 | 55.8 | |
| *fzo-1(mut);pdr-1(mut);ΔmtDNA* | 18 | 9 | |

**Table S2: Coverage, haplotyping and summary statistics**

| Patient | Haplogroup | Read coverage (Average ± SD) | Covered positions (>500X) |
|---|---|---|---|
| K_V13 | H1a | 6169.76 ± 623.32 | 16560 |
| K_V14 | H1a | 6201.94 ± 596.07 | 16560 |
| K_V17 | H1a | 5882.87 ± 636.84 | 16560 |
| K_VI9 | H1ab | 6119.97 ± 632.06 | 16555 |
| Z_0106 | H1ae1 | 6233.64 ± 592.38 | 16560 |
| Z_1007* | NA | 6711.62 ± 1374.66 | 952 |
| Z_1008 | H1ae1 | 6165.76 ± 616.65 | 16560 |
| N_SD1 | T2 | 6036.13 ± 562.88 | 16558 |
| N_SD2 | T2 | 5806.99 ± 945.31 | 16543 |
| N_SD3 | T2 | 5674.63 ± 761.54 | 16560 |

*Only for mtDNA positions 13607-14578

### Table S3: Heteroplasmy in analyzed patient samples

| Sample | mtDNA position | Heteroplasmy level | Best reads | Secondary reads |
|---|---|---|---|---|
| **K_V13** | 16172 | 6.24 | T | C |
| | 310 | 1.59 | T | C |
| | 15880 | 1.19 | A | G |
| **K_V14** | 6899 | 10.59 | G | A |
| | 16172 | 10.08 | T | C |
| | 3447 | 1.91 | A | C |
| | 4665 | 1.88 | G | A |
| | 310 | 1.29 | T | C |
| **K_V17** | 16172 | 10.34 | C | T |
| | 6899 | 8.82 | G | A |
| | 310 | 1.06 | T | C |
| **K_VI9** | 152 | 9.93 | T | C |
| | 3492 | 4.26 | A | C |
| | 15515 | 2.94 | A | G |
| | 378 | 2.47 | C | T |
| | 3447 | 2.30 | A | C |
| | 3677 | 2.13 | A | G |
| | 310 | 1.60 | T | C |
| | 6764 | 1.26 | G | A |
| | 14882 | 1.23 | A | G |
| | 3475 | 1.17 | A | C |
| **Z_1007** | 13830 | 20.28 | T | C |
| | 13711 | 2.86 | G | A |
| | 13869 | 2.82 | T | C |
| | 13855 | 2.78 | C | T |
| | 13707 | 2.58 | G | A |
| | 14053 | 2.37 | A | G |
| | 13899 | 2.23 | T | C |
| | 14034 | 2.13 | T | C |
| | 13708 | 2.08 | G | A |
| | 14088 | 2.07 | T | C |
| | 13712 | 2.06 | C | T |
| | 14041 | 1.92 | C | T |
| | 13833 | 1.91 | A | G |
| | 14020 | 1.75 | T | A |
| | 13674 | 1.74 | T | C |
| | 14016 | 1.71 | G | A |
| | 14033 | 1.69 | T | C |
| | 14073 | 1.65 | C | T |
| | 14048 | 1.60 | T | C |
| | 14122 | 1.56 | A | C |
| | 13908 | 1.48 | C | T |
| | 13905 | 1.45 | C | T |
| | 14133 | 1.42 | A | C |
| | 13686 | 1.37 | A | G |
| | 13929 | 1.33 | C | T |
| | 13934 | 1.31 | C | T |
| | 13698 | 1.31 | T | C |
| | 13811 | 1.29 | C | G |
| | 13920 | 1.25 | C | T |
| | 13809 | 1.24 | C | A |
| | 14305 | 1.23 | G | A |
| | 13680 | 1.21 | C | T |
| | 13968 | 1.15 | G | A |
| | 14280 | 1.15 | A | G |
| | 14287 | 1.13 | T | C |
| | 13691 | 1.12 | A | G |
| | 13950 | 1.07 | C | T |
| | 14275 | 1.06 | C | T |
| | 14276 | 1.01 | C | G |
| | 13945 | 1.00 | A | G |
| **Z_1008** | 13830 | 35.00 | T | C |
| | 9722 | 2.16 | T | C |
| | 5368 | 1.22 | C | G |
| **Z_0106** | 13830 | 25.38 | C | T |
| | 310 | 1.08 | T | C |
| **N_SD1** | 4491 | 1.80 | G | A |
| **N_SD2** | 11963 | 1.37 | G | A |
| | 11812 | 1.31 | G | A |
| | 12007 | 1.29 | G | A |
| | 11914 | 1.22 | G | A |
| | 12013 | 1.14 | A | G |
| | 11887 | 1.02 | G | A |
| **N_SD3** | 5070 | 1.54 | A | G |

**Table S4: A list of *C. elegans* strains used in this study**

| Strain | Abbreviation | Nuclear genotype | Mitochondrial genotype |
|---|---|---|---|
| **N2** | wild type (WT) | --- | -- |
| **AM134** | Q0 | *unc-54p::Q0::YFP* | -- |
| **ABZ271*** | *fzo-1* | *fzo-1(tm1133)*<br>derived from CU5991 | -- |
| **ABZ283*** | *pdr-1* | *pdr-1(gk448)*<br>derived from VC1024 | -- |
| **ABZ270*** | ΔmtDNA | -- | uaDf5/+<br>derived from LB138 |
| | G1ht | *fzo-1 +/-* | uaDf5/+ |
| **ABZ272** | *fzo-1(mut);ΔmtDNA* | *fzo-1(tm1133)* | uaDf5/+** |
| **ABZ273** | *fzo-1(wt);ΔmtDNA* | *fzo-1(+)* | uaDf5/+ |
| **ABZ274** | G4mut->G6wt | *fzo-1(+)* | uaDf5/+** |
| **ABZ275*** | 1kbΔmtDNA | -- | *bguDf1*/+<br>derived from VC41028 |
| | G1ht 1kbΔmtDNA | *fzo-1 +/-* | *bguDf1*/+ |
| **ABZ276** | *fzo-1(mut);1kbΔmtDNA* | *fzo-1(tm1133)* | *bguDf1*/+** |
| **ABZ277** | *fzo-1(wt);1kbΔmtDNA* | *fzo-1(+)* | *bguDf1*/+ |
| **ABZ278** | G4mut->G6wt | *fzo-1(+)* | *bguDf1*/+** |
| **ABZ279*** | 4kbΔmtDNA | -- | *bguDf2*/+<br>derived from VC20469 |
| | G1ht 4kbΔmtDNA | *fzo-1 +/-* | *bguDf2*/+ |
| **ABZ280** | *fzo-1(mut);4kbΔmtDNA* | *fzo-1(tm1133)* | *bguDf2*/+** |
| **ABZ281** | *fzo-1(wt);4kbΔmtDNA* | *fzo-1(+)* | *bguDf2*/+ |
| **ABZ282** | G4mut->G6wt | *fzo-1(+)* | *bguDf2*/+** |
| **ABZ285** | *pdr-1;*ΔmtDNA | *pdr-1(gk448)* | uaDf5/+ |
| **ABZ284** | *fzo-1;pdr-1* | *fzo-1(tm1133);pdr-1(gk448)* | -- |
| **ABZ286** | *fzo-1(mut);pdr-1(mut); ΔmtDNA* | *fzo-1(tm1133);pdr-1(gk448)* | uaDf5/+** |
| **ABZ287** | *fzo-1(wt);pdr-1(mut); ΔmtDNA* | *fzo-1(+);pdr-1(gk448)* | uaDf5/+ |

* Strains were outcrossed with our lab N2 stock at least four times
** ΔmtDNA levels were lost over generations

**Table S5**: **Primers used for *C. elegans* experiments**

| Primer | Primer sequence |
|---|---|
| *fzo-1* | F-TTCCTGCATCCGGTCTCATT |
| | R-AGTCGGCATTCCCCTGATTC |
| *pdr-1* | F-CGGTCGCTGTGAGTTTAGAA |
| | R-GGAGTACAGCATTCTTCGCA - |
| ΔmtDNA | F -TGAGACTTTTAATTATTTACATCCC |
| | R-CAGTGCATTGACCTAGTCATC- |
| 1kbΔmtDNA | F-ATACTTTACCATTAAGGTCAGTAATTTCTA |
| | R- GTTGTTTCTCAATTAATAAAATTATAACCCC |
| 4kbΔmtDNA | F-TCAAGGAGGATTGGCAGTTTGA |
| | R-ACCTCTAAAAACCGATAAACCAAAA |
| wt of ΔmtDNA and total of 4kbΔmtDNA | F -ATGGGATGTTGGTGACATTGC |
| | R-TGCTATTAACCTATCGGGCGTA |
| wt of 1kbΔmtDNA | F-ATACTTTACCATTAAGGTCAGTAATTTCTA |
| | R-TCACGCTACAGCAGCATAAAC |
| wt of 4kbΔmtDNA | F-TCAAGGAGGATTGGCAGTTTGA |
| | R-TCTAGTACCAACCATACCAGATCA |
| Total of ΔmtDNA and 1kbΔmtDNA | F-TCGGTGGTTTTGGTAACTGAT |
| | R-CTGTATTCTCCGGATTACGAG |
| Genomic DNA | F-CTGGAAGAAGATAATTATTTTCC |
| | R-CTGTATTCTCCGGATTACGAG |

**Table S6: Primers used for Sanger sequencing of human mtDNA fragments**

| Primer | mtDNA Positions | DNA sequence |
|---|---|---|
| Forward primer | (NC_012920) | CACCATTAGCACCCAAAGCT |
| Reverse primer | 15997-16016 | TGATTTCACGGAGGATGGTG |
| Forward primer | 16420-16439 | TCGAATAATTCTTCTCACCC |
| Reverse primer | 13607-13627 | GATTGTTAGCGGTGTGGTCG |
